# Nest records, Nest Site selection of *Gyps bengalensis* White-rumped Vulture and the role of Feeding Station in Kangra, Himachal Pradesh

**DOI:** 10.1101/2020.07.29.218362

**Authors:** Archi Sehgal, Krishan Kumar, Rubina Rajan, Upmanyu Hore

**Author notes:** Corresponding author Dr. Upmanyu Hore, Assistant Professor, Amity Institute of Forestry and Wildlife, Noida,Uttar Pradesh, India, Email ID.

## Abstract

Active nests of *Gyps bengalensis* White-rumped Vulture depends on elevation and aspects for nest site selection, while feeding station plays a significant role for determining the position of the nesting sites.This study attempted to record nest count for the breeding period 2018, identify key variables for the nest site selection and understand the role of feeding station in the nest site selection. Nest counts were conducted during the breeding period, each nest was categorized into active and inactive nest based on assesment of different components. Simultaneously, different variables (tree height, tree species, elevation and aspect) were recorded for each nest. Aerial distance was used to determine the role of feeding station for selecting the nesting sites. From the 24 nesting sites, 352 active nests were recorded, and a significant Pearson’s correlation for elevation and aspect were drawn. We found, vulture prefer single tree species for nesting. We also found that, 71% (n = 17) nesting sites located within radial distance of 20 km of the feeding station. High congregration of active nests within short radial distance from feeding station, signify the positive impact of management of feeding station by the wildlife wing of Forest department since 2008, for the ex-situ conservation of critically endangered *Gyps bengalensis* White-rumped Vulture

## Introduction

Among 23 species of vultures, 12 species are currently classified as “Near Threatened” or “Endangered” (IUCN 2017). In India, 106 Raptor species are found which makes 18 percent of 572 species that are spread all over the world (Ali et al. 1987, Naoroji & Schmitt 2007). Of these, eight species of vultures are reported from Himachal Pradesh: *Gyps bengalensis* White-rumped Vulture, *Gyps himalayensis* Himalayan Griffon *Gyps fulvus* Eurasian Griffon, *Gypaetus barbatu* Lammergeyer, *Gyps indicus* Indian Long-Billed Vulture, *Aegypius monachu* Cinereous Vulture, *Neophron percnopterus* Egyptian Vulture, and *Sarcogyps calvus* Red Headed Vulture (Puri et al. 2018). 30 years ago, the vulture population in Northern-Central India, appeared to be highest in urban areas (Galushin 1971). The evidence of catastrophic decline reported during the late 1990’s from Keoladeo National park, Rajasthan (Prakash 1999). Various hypotheses were put forward to investigate the cause for this rapid decline (Pain et al. 2003, Prakash et al. 2003, Oaks et al. 2004). Later it was discovered, a non-steroidal anti-inflammatory drug (NSAID): diclofenac was the main causative factor for rapid population decline across the Indian subcontinent (Oaks et al. 2004). White-rumped Vulture is one of four Critically Endangered species of vultures in India, others are Indian Long-billed Vulture, Slender-billed Vulture Red-headed Vulture (Prakash 1999, Green et al. 2007, Prakash et al. 2007)

White-rumped Vulture is the smallest among Gyps species, feed exclusively on carrion, they are gregarious and social, they forage in flocks, roost and nest in colonies (Sharma 1970). They breed and nest from October–April. Pairs are monogamous, raise one nestling per year, the incubation period is approximately 50-56 days, the nestling period lasts approximately 104 days (Gillbert et al. 2002, Thakur 2015, Baral et al. 2005). This species is known to show partial migration that ranges from 1000-2000 square kilometres and sometimes across the borders between Nepal and India (Clements et al. 2013, Ferguson-lees 2001).

Their population has drastically been affected in the last decade as a result of several factors -the use of diclofenac, shortage of food and electrocution. Thus, to conserve the breeding population, it is important to safeguard their nesting habitats which vary in spatial scale and it has been reported for other vulture species that the establishment of feeding stations can significantly increase the survival of first-year vultures (Piper et al. 1999) and can also affect the growth in nesting colonies as seen in California Condor (Snyder & Snyder 2005). The present study aimed to identify nesting sites of White-rumped Vulture and determine nest site variables such as aspect of the nest, elevation, the height of nesting tree. Also, to investigate the role of feeding stations, which has been established as a species-specific conservation measure by Forest department of Himachal Pradesh.

## Materials and Methods

### Study Area

This study was carried out in the Kangra district of Himachal Pradesh. This district is traversed by varying altitudes of Shivaliks, Dhauladhar and Himalayas from northwest to southeast. The altitude varies from 500 m above mean sea level to around 5000 m above the mean sea level. The climate of the district varies from sub-tropical in lower hills and valleys to sub-humid in the mid-hills and gets temperate in high hills. The forest cover of Kangra district includes 297 km^2^area of very dense forest and 1279 km^2^ of moderate dense forest (FSI 2017). We carried out nest survey within the boundary of our study area with the help of Forest staff (Fig.1). We also relied on the knowledge and experience of the locals for tracing nests. We visited the feeding station located in Nagrota Surian village of Kangra district, constructed in an open land area of 100 metres x 100 metres. It is fenced to exclude facultative scavengers and maintained by Forest department since 2008. The dead livestock comes from nearby villages, which is transported by members of a particular community called ‘MOOCHIS’ to the feeding station.

**Figure 1.**
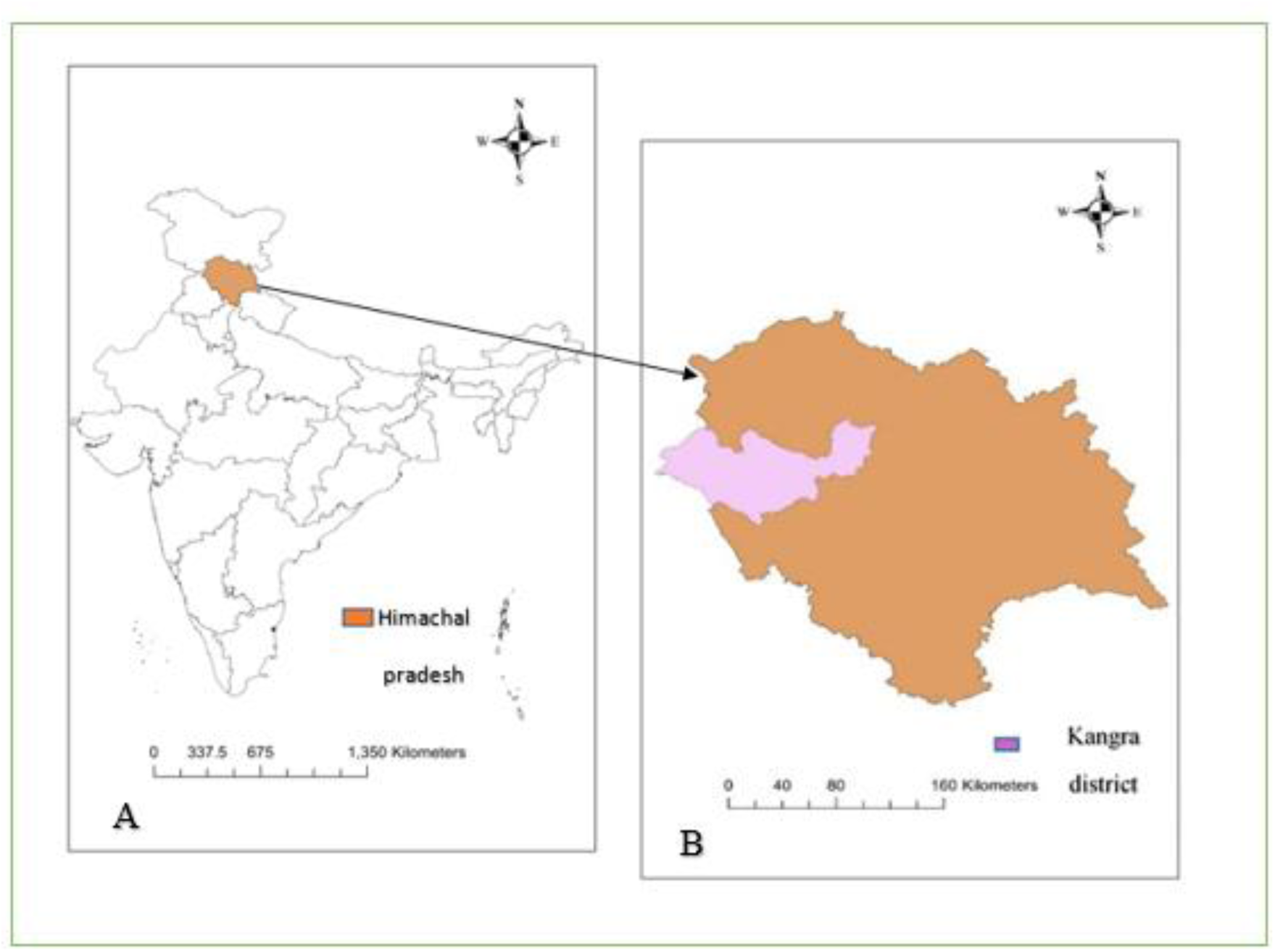
Map of study area: (A) highlights the state of Himachal Pradesh, (B) marks the Kangra district within Himachal Pradesh.

## Data collection

### Nest records

The nest survey was carried out during the hatching and fledgling period of White-rumped Vulture from January 2018 to April 2018 with the help of forest department. The prior reports of forest department recorded a total 48 nesting sites in the same district, some of these nesting sites were abandoned by White-rumped Vulture over the years, we scanned all the 48 nesting sites which were previously identified and regularly monitored during the breeding season in different beats and ranges of forest in Kangra. We considered those nests with the presence of parent, fresh droppings and the fledgling as “active nest”, Whereas nest with no parent, fledglings or fresh droppings were identified as “inactive nest” (Chomba et al., 2013).Each of the nests were observed using Olympus binocular 50 × 100 from a distance to identify it as an “active” or “inactive” nest. We considered “beat” as a unit for the nesting site, the beats having at least one active nest were recorded as “active nesting site”. While beats with zero active nests were recorded as” inactive nesting site”. Each nesting site was extensively surveyed on foot, observations were recorded for each nest. To avoid duplication, we followed unidirectional movement for the survey.

### Nest site variable

At each nest we recorded the various variables (Table 1) to determine the selection of nesting sites. We selected some important variables based on literature (Thakur and Narang 2012, Baral et al. 2005, Freund et al. 2017). We recorded GPS coordinates, the height of nesting tree, elevation and aspect of the nest. To determine the height of nesting tree a clinometer was used. To record the elevation, Garmin eTrex GPS handset was used to take readings for each of the nests and for aspect, we used DEM map in ArcGIS 10.3. using each nest’s GPS coordinates. We also recorded tree species on which the nest was build.

**Table 1.** Description of Nest variables

| Nest site variables | Measurement Unit |
| --- | --- |
| Tree species | At species level |
| Height of each nesting tree | Measured in metres(m) |
| Elevation of each nest | Measured in metres(m) |
| Aspect of nest | Measured in degree (North- 0°, East- 90°, South-180°, West-270°) |

### Feeding Station

To determine the role of feeding station in the selection of nesting sites by White-rumped Vulture, we used the Garmin eTrex GPS handset to record GPS location of feeding station and each nesting site. The aerial distance was calculated between the feeding station and each active nesting site using Google Earth Pro. Also, we calculated the number of nesting sites falling in radial distance of 10 KM, 20 Km, and 30 Km through the same method. We tried to establish link between the ratio of inactive nest to active nest at each nesting site with the distance from the feeding station.

### Statistical Analysis

We listed all active and inactive nest found at each nesting site. Nesting sites are classified as “active nesting site” and “inactive nesting site” based on the number of active nests recorded within it. We used Pearson’s correlation to identify variables that are correlated with the nesting site selection. We consider each nest as a sample unit for the analysis. This study also looked at the role of feeding station in the nesting site selection. Thus, each nesting site were taken as unit of analysis for the spearman’s rank correlation, the ratio of inactive to active nest at each nesting was correlated with the distance to the feeding station (Vlachos 1998).

## Results

### Nest record

Total 387 nests of Critically Endangered White-rumped Vulture were recorded of which 352 were active with fledglings. We recorded 24 active nesting sites having at least one active White-rumped Vulture nest (Fig.2). From the (Table 2), the lowest number of nests was recorded from ‘Bassa’ (n=2), situated next to the roadside. While the highest number of nests recorded at ‘Pathiar’ site (n= 43), located in a dense forest area.

**Figure 2:**
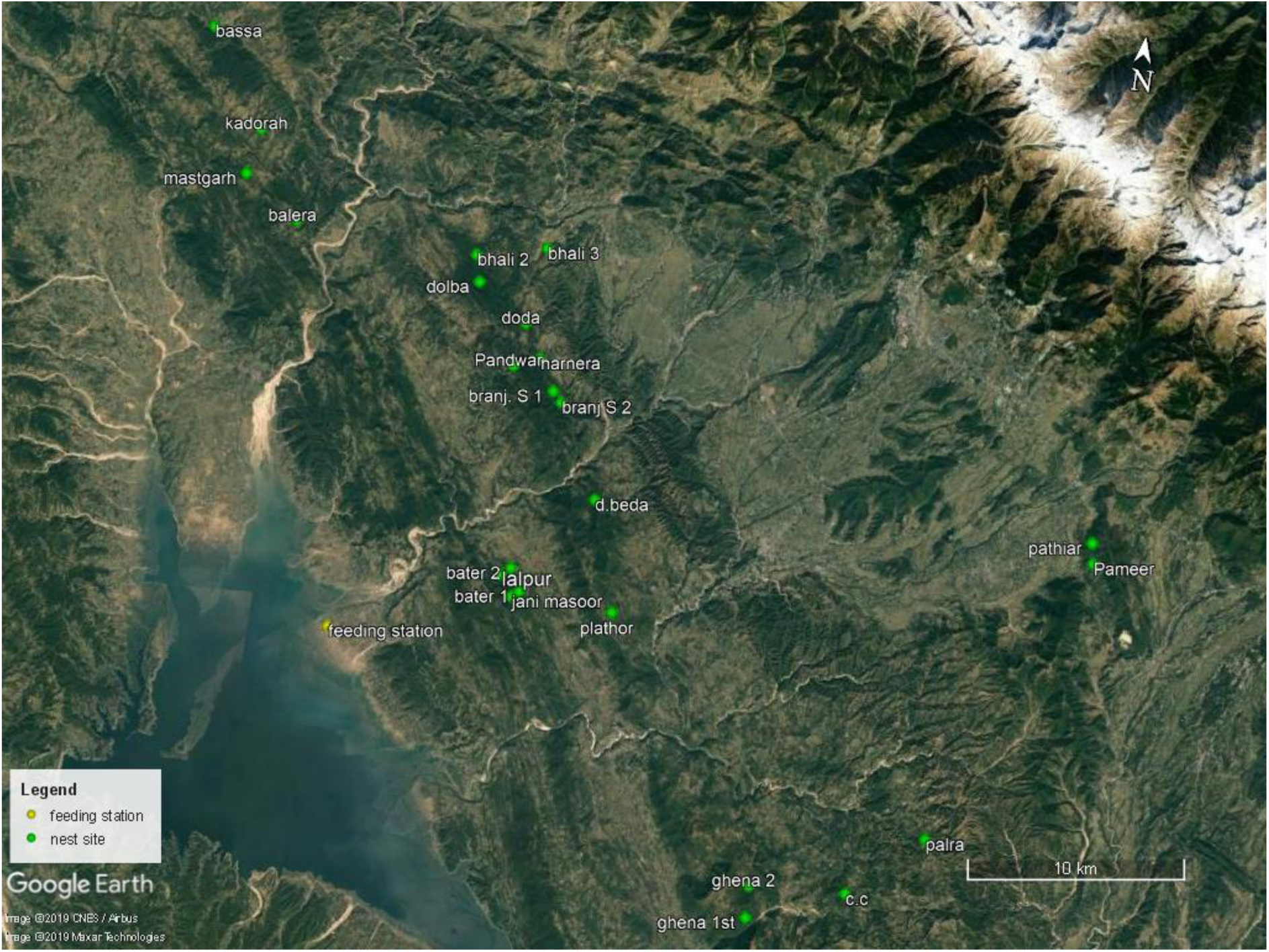
GPS location of 24 active nesting sites of White-rumped Vulture, marked with green dots and Feeding station highlighted with yellow dot.

**Table 2.** Description of 24 active nesting sites with total nest and active nest with fledgling found at each site, also the distance between nesting site and feeding station is given below:

| **Nesting site** | **Name of Nesting site** | **Total number of nests recorded** | **Number of fledglings** | **Aerial distance with respect to Feeding station (km)** |
| --- | --- | --- | --- | --- |
| **1** | Bater 1<sup>st</sup> | 19 | 16 | 8.94 |
| **2** | Bater 2<sup>st</sup> | 6 | 4 | 8.92 |
| **3** | Jani Masoor | 30 | 27 | 8.55 |
| **4** | Branj s. 1<sup>st</sup> | 15 | 13 | 14.96 |
| **5** | Branj s. 2<sup>nd</sup> | 17 | 15 | 14.94 |
| **6** | Harnera | 4 | 4 | 15.68 |
| **7** | Pandwar | 24 | 22 | 14.72 |
| **8** | C.C | 26 | 24 | 26.78 |
| **9** | Pameer | 7 | 7 | 35.3 |
| **10** | Bhali 2 | 6 | 6 | 18.39 |
| **11** | Mastgarh | 13 | 12 | 21.37 |
| **12** | Kadroah | 30 | 26 | 23.21 |
| **13** | Lalpur | 40 | 38 | 8.33 |
| **14** | Doda | 5 | 4 | 16.61 |
| **15** | Palra | 15 | 14 | 29.21 |
| **16** | Ghena 1st | 11 | 10 | 23.41 |
| **17** | Pathiar | 45 | 43 | 35.28 |
| **18** | Bhali 3<sup>rd</sup> | 16 | 15 | 20.3 |
| **19** | Dolba | 25 | 22 | 17.22 |
| **20** | Balera | 9 | 9 | 18.81 |
| **21** | Gheena 2 | 3 | 3 | 22.73 |
| **22** | Bassa | 2 | 2 | 28.21 |
| 23 | D. beda | 15 | 12 | 13.61 |
| 24 | plathore | 4 | 4 | 13.9 |

### Nest Site Variable

We found a significant correlation between elevation and nest i.e., nest site with average elevation 671 ± 68 m had a greater number of nests within nesting site (Pearson’s correlation, r= 0.526, *P <* .0001). When calculating correlation between aspect and nests: we found a significant correlation between aspect and nests (Pearson’s correlation, r = .154, *P <* .0007). White-rumped Vulture prefers nesting site more towards the south (22%, n= 78), south-west (17%, n=60) and north (18%, n=68) and few nesting colonies in east (5% n=18), north-east (8% n= 29), northwest (4% n= 17) and west (12% n= 43) direction. No significant correlation was found between height of nesting tree and nest site (Pearson’s correlation, r= -.10, P < .06). We found that all nests of White-rumped Vulture were built on pine trees (*Pinus roxburghii*) and the average nesting tree height was 20.8 ± 2.7 m.

### Feeding station

On calculating the correlation between ratio of inactive and active nest at each site to the distance of feeding station we found, significant correlation (Spearman’s correlation r = 0.428, *P* <0.037), we observed (Fig.3) that with the increase in distance the ratio of inactive to active increases, thus nesting sites are confined to a particular distance from the feeding station. 71% (n=17) of nesting sites were recorded within the radial distance of 20 Km from the feeding station. We found all the nesting sites are located with an average aerial distance of 19.14 ± Km from the feeding station.

**Figure 3.**
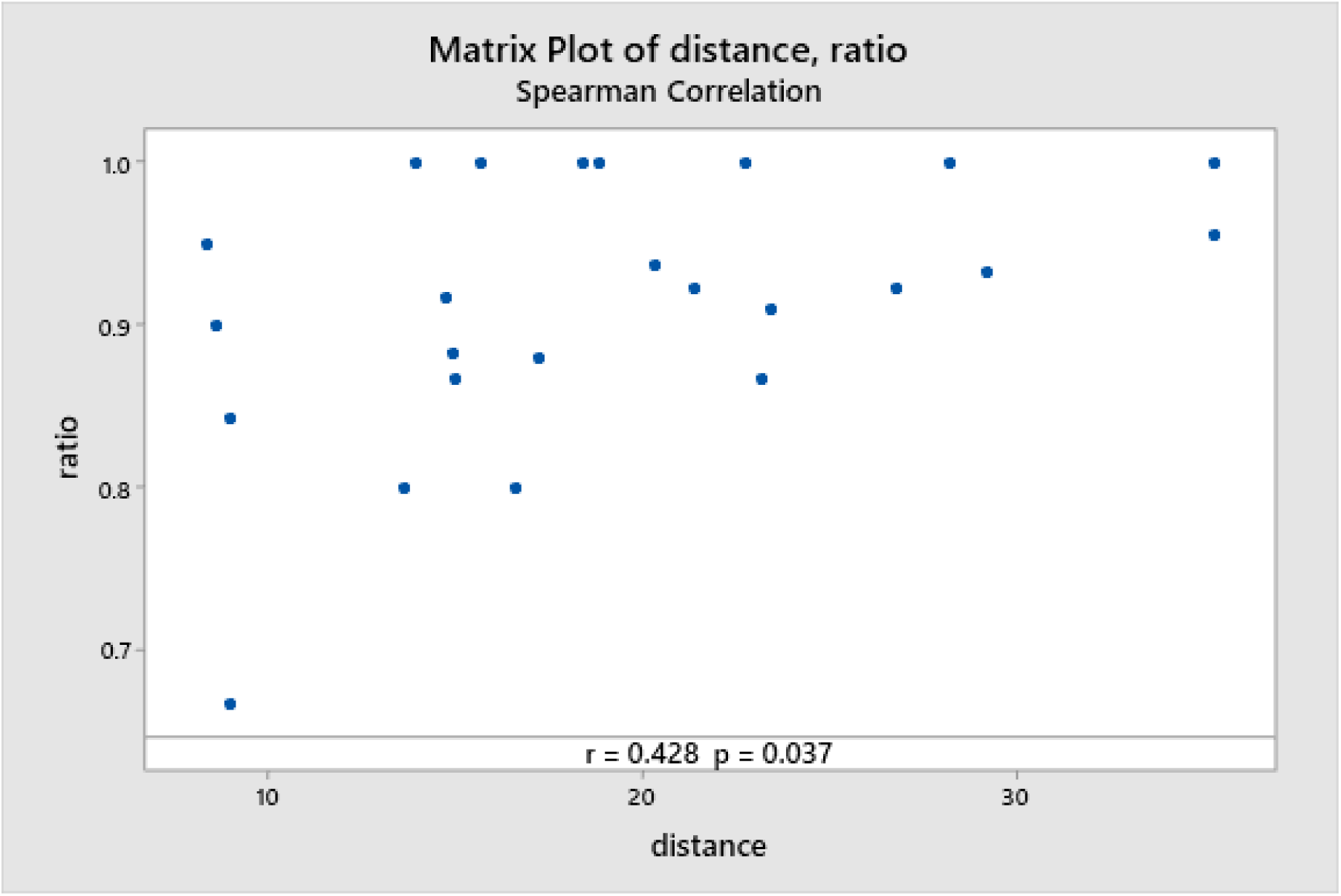
Scatter plot showing spearman’s rank correlation between ration of inactive and active nests at each nesting site to the aerial distance of Feeding station

## Discussion

### Nest record

The present study area that sustains 352 active White-rumped Vulture nests for the breeding season of 2018-19 indicates that Kangra district of Himachal Pradesh is an important conservation site. In a study done during the breeding season of 2009-2011, 19 active nesting sites of White-rumped Vulture were reported within the present study area (Thakur and Kataria, 2012). It was assumed that some critical ecological factors may support the breeding population in the low laying Shivaliks hills of Kangra district. This may also be attributed to the awareness among the people regarding the dangers of diclofenac drug and accompanied by the low usage of diclofenac around nesting sites (Thakur and Kataria, 2012).

### Nest Site Variable

The results of this study showed that the selection of the nesting site is influenced by key variables. The two most important variables that significantly contribute to the nesting site selection are **elevation** and **aspect**. The height of nesting tree was not found to play a role in nesting site selection as the preference for nesting tree height varied greatly which is consistent with other studies. White-rumped Vulture was reported to build nest as low as four to five metres where large trees are absent (Naroji 2007). Similar observations were reported from Gujarat with an average tree height of 16.4 m (Gadhvi 2006). Ramakrishan 2014, reported that nests at a height between 18 to 36 metres in Segur plateau, Tamil Nadu. Thus, indicating that the White-rumped Vultures are quite flexible with respect to the height of the nesting tree and tree species according to the topography and abundance of the tree species present in the area. The variable which has shown significant relationship with the selection of the nesting site is elevation, with an average elevation of 671 ± 68 metres. The White-rumped Vulture are known to restrict their nesting to low-land areas (Naroji 2007, Giri et.al. 2002). *Pinus Roxburghii* mostly occur in lower Shiwaliks of Kangra district between the elevation of 450 metres to 1050 metres. White-rumped Vulture may be using these pine trees at this elevation due to the absence of other dominant tree species with the average height of 20.8 metres in the area. The results indicate that the selection of the nesting site is also dependent on the aspect of the nest. Most nests were observed to be oriented in the north and south direction. In a similar study done on the same species in Moyar valley of Tamil Nadu it was found that west, north and east orientated nests were in the majority (Venkitachalam and Senthilnathan, 2018). Other species such as *Gyps Indicus* Indian Long-billed vulture and Gyps *fulvus* Eurasian Griffon were also reported nesting in south-east and south oriented nest. This indicates that aspect plays a role in nest building. Orientation may be instrumental in protecting them from inclement conditions like directional movement of cold winds or direct solar radiation and facilitates them to raise their body temperature after nocturnal dip as temperature is key to reproductive success during early breeding stages. (Xirouchakis and Mylonas, 2013; Venkitachalam and Senthilnathan, 2015).

### Feeding station

Many studies support the impact of feeding stations in facilitating recolonization of scavenging raptors (Mundy et al. 1992, Oro et al. 2008, Lieury et al. 2015). In our study, all nesting sites were located at an average aerial distance of 19.14 ± 7.78 kilometres. The presence of active nests shows a strong correlation with the distance from the feeding station. The possible reasons for the correlation could be reduction in the foraging time due to proximity to the feeding station and also the availability of food at feeding stations which reduces the risk of mortality (Snyder and Snyder, 2005). Feeding stations can play a significant role in subsequent slight population recovery, as reported in long-billed vulture (Gilbert et al., 2007, Balmford, 2013) and for California Condor (Snyder and Snyder, 2005) also it improves the reproductive success and survival of first year juveniles (Piper et al., 1999, Grande et al., 2009, Margalida and Colomer, 2012). Apart from the feeding station, the present study area also supports a high livestock population: 8-9 lakh (Sharma and Shilpa, 2016), and the carcasses of these livestock are usually dumped near the roadside by the locals (Thakur and Kataria, 2012). This practice is common where the feeding stations are far from the respective locations or absent. As predicted, proximity to concentrated food resources favors the establishment of breeding colonies (Lieury et al., 2015). Thus, we reported a similar phenomenon with 71% of nesting sites of White-rumped Vulture were located within 20 km of radius of the feeding station.

Vultures are an integral part of the terrestrial ecosystem and their loss can translate into a grave crisis. (Verner et al., 1986). Their population and nesting success are dependent on the availability of nesting sites (Newton,1979). The data on nesting success of a species is of conservation significance to understand the nature and extent of habitat use and decline of the species (Misher, 2017). Use of Diclofenac drug and the retaliatory killings of carnivores using poison bait are the major threats to scavengers like White-rumped Vultures. In this scenario feeding stations that provide safe food resources for the vultures is an important conservation step shown to be successful in many cases. Feeding stations have shown to increase breeding propensity (Oppel, 2016). Proper monitoring and management are required in the regulation of feeding stations in order to get the desired effects. It also plays a role to minimize negative effects such as loss of scavenging service efficiency and habituation to feeding stations. The above can be controlled by regulating the number of feeding stations and the frequency of supply to these feeding stations. Feeding stations can also be a platform for creating awareness about these creatures, improve sanitation and also boost tourism as seen in the case of vulture restaurants of Nepal.

Such a study indicates that adaptive management in which active management and research occurs simultaneously is a useful strategy to maximize management efficiency. As a large amount of human and financial resources are being spent on the conservation actions aimed at halting the decline of species worldwide, there is a need to evaluate the conservation actions and its success.

## Acknowledgments

Special thanks to H.P. Forest Department (Wildlife wing) for all logistics and support. We thank the Department of Forestry and Wildlife, Amity University for providing us technical support throughout the study.

## Notes

### Competing Interest Statement

The authors have declared no competing interest.

